# Geometric regulation of histone state directs melanoma reprogramming

**DOI:** 10.1101/872226

**Authors:** Junmin Lee, Thomas G. Molley, Christopher H. Seward, Amr A. Abdeen, Huimin Zhang, Xiaochun Wang, Hetvi Gandhi, Jia-Lin Yang, Katharina Gaus, Kristopher A. Kilian

## Abstract

Malignant melanoma displays a high degree of cellular plasticity during disease progression, making classification of the heterogeneous population and selection of an appropriate therapy challenging. Signals in the tumor microenvironment are believed to influence melanoma plasticity through changes in the epigenetic state to guide dynamic differentiation and de-differentiation events that underlie tumorigenicity and dissemination. Here we uncover a relationship between geometric features at perimeter regions of multicellular melanoma aggregates, and reprogramming to a stem cell-like melanoma initiating cell (MIC) through histone marks H3K4Me2 and H3K9Ac. Using an in vitro tumor microengineering approach, we find concurrent expression of molecular MIC markers and spatial enrichment of these histone modifications at perimeter features. Chromatin immunoprecipitation and sequencing analysis demonstrates broad regulation of genes associated with SOX-, ETS-, and USF-families. SOX10 and PRDM14, transcriptional regulators with a putative role in several cancers, overlap with H3K9Ac and show elevated expression in cells along regions of perimeter curvature. siRNA knockdown of the epigenetic modifier PRDM14 abolishes the MIC phenotype suggesting a role in regulating melanoma heterogeneity. Our results suggest mechanotransduction at the periphery of melanoma tumors may orchestrate the activity of epigenetic modifiers to regulate histone state, cellular plasticity, and tumorigenicity.

## 1. Introduction

Malignant transformation and metastatic spread are known to be mediated by both genetic abnormalities^[1]^ and epigenetic alterations.^[2]^ Epigenetics, defined as heritable change in gene expression occurring independent of changes in primary DNA sequence, is strongly implicated in the underlying mechanisms of cancer progression.^[3]^ Microenvironment-mediated epigenetic regulation of cancer-related gene expression through DNA methylation, histone modification, and chromatin compartments is now believed to take part in a broad spectrum of cancer behaviors ranging from initiation to phenotypic alteration.^[4]^ Histone modifications, including methylation, phosphorylation, and acetylation are covalent post-translational modifications to histone proteins. These modifications allow histones to alter the structure of chromatin, resulting in transcriptional activation or repression, that affect changes in cell behavior. For example, histone H3 lysine 4 di/tri-methylation (H3K4me2/3) and histone H3 acetylation (H3ac) are generally associated with gene activation,^[5]^ whereas H3K27me, which marks active cis-regulatory elements, is associated with gene inactivation.^[6]^ The detection of cancer-specific changes through histone modifications as epigenetic biomarkers has potential for clinical prediction, diagnosis, and therapeutic development.

Malignant melanoma-initiating cells (MICs), also referred to as melanoma repopulating cells or melanoma stem cells, are a dynamic sub-populations of cells that may arise during progression with tumor initiating capacities.^[7]^ Unlike the clonal evolution model describing how a single cell accumulates genetic and epigenetic changes until becoming a cancer tumor cell,^[1]^ the cancer stem cell model suggests a hierarchical organization (unidirectional) of cancer cells, according to their tumorigenic potential that has important implications for cancer therapy with stem cell-specific treatment regimens.^[8]^ However, accumulating evidence surrounding cancer plasticity supports a new emerging model of tumorigenicity, in which dynamic plasticity facilitates malignant cells to revert to a stem cell-like phenotype.^[9]^ Recently, we and other groups have shown that cancer cells are more plastic than previously anticipated. For instance, conversion to a stem cell-like state has been guided by microenvironment-mediated epigenetic regulation of gene expression including factors such as pH,^[10]^ radiation,^[11]^ stiffness,^[12]^ hypoxia,^[13]^ and interfacial stress.^[14]^ These microenvironment parameters are not mutually exclusive and likely integrate in a context-dependent fashion during progression to guide tumor heterogeneity underlying progression. Hence, we hypothesize that if tumor cells are put into a specific context which facilitates reprogramming to the MIC phenotype, specific histone modifications may be used to understand the mechanisms underlying phenotypic alterations during progression. Our use of microengineering based on soft lithography allows us mimic aspects of the tumor microenvironment, thus effectively deconstructing the biophysical cues of stiffness and geometry to probe how these parameters provide a context to facilitate epigenetic reprogramming to a stem cell-like tumorigenic state.

## 2. Results and Discussion

### 2.1. Geometric cues regulate histone methylation and acetylation in melanoma

To classify histones linked to epigenetic reprogramming from melanoma to the MIC state, we employed microengineered hydrogels that we previously demonstrated will coordinate enhancement of the MIC phenotype with spatial control (Fig. S1). In our previous study, B16F0 murine melanoma cells cultured for five days expressed higher levels of MIC markers at the periphery of microaggregates, with stem cell-like characteristics demonstrated in vitro and in vivo.^[14]^ To understand how geometric cues at the perimeter of microaggregates of melanoma cells will influence histone state we characterized a panel of histone marks that are implicated in controlling oncogenic gene activation. We first analyzed methylation state at Histone H3; histone H3 lysine 4 methylations (mono, di, and tri) were studied because these are known as active histone marks.^[5]^ In addition, Jarid1B (gene name: KDM5B) was also investigated because it is a common molecular marker of MICs, and is the histone lysine demethylase for H3K4me3/2/1 with pronounced roles in different cancer types.^[15]^ For example, overexpression of Jarid1B in the MDA-MB 231 breast cancer cells suppressed malignant characteristics such as cell migration and invasion,^[16]^ while overexpression of Jarid1B in melanoma^[17]^ or immortalized normal breast cancer cells (MCF10A)^[18]^ was found to enhance metastatic progression or cell invasion, respectively. Another representative histone mark associated with transcriptional activation, histone H3 lysine 36 methylation (H3K36me2), and a histone mark correlated with transcriptional repression, histone H3 lysine 9 methylation (H3K9me3), were also employed.

We cultured cells for five days on five different micropatterned hydrogel substrates with the same area (50,000 μm^2^) or on non-patterned protein-stamped hydrogel substrates (10 kPa gels for both patterned and non-patterned) and immunostained for histone methylation state (Fig. 1A, S2, and S3). Interestingly, H3K4me2 and H3K36me2 expression co-localized with cells adopting the MIC phenotype at the periphery of microaggregates. We selected cells cultured for five days in the spiral shape for flow cytometry analysis because we previously found that this shape will augment the MIC phenotype^[19]^ through high interfacial boundary (perimeter/area) and high curvature.^[14]^ Similar to the immunofluorescence results, cells cultured in the spiral patterns display higher levels of H3K4me2 and H3K36me2 expression compared to those cultured on non-patterned surfaces (Fig. 1B). To gain an understanding into the mechanism underlying the observed spatial distribution of histone marks, cells were grown in circular shapes, followed by quantification of histone marks in two different regions (outside and inside) within the same area. Cells cultured at the perimeter displayed significantly higher levels of H3K4me2 compared to those cultured at central regions (Fig. 1C), suggesting that regulation of gene expression associated with reprogramming of melanoma cells into the MIC phenotype could be linked to the H3K4me2 mark.

**Figure 1.**
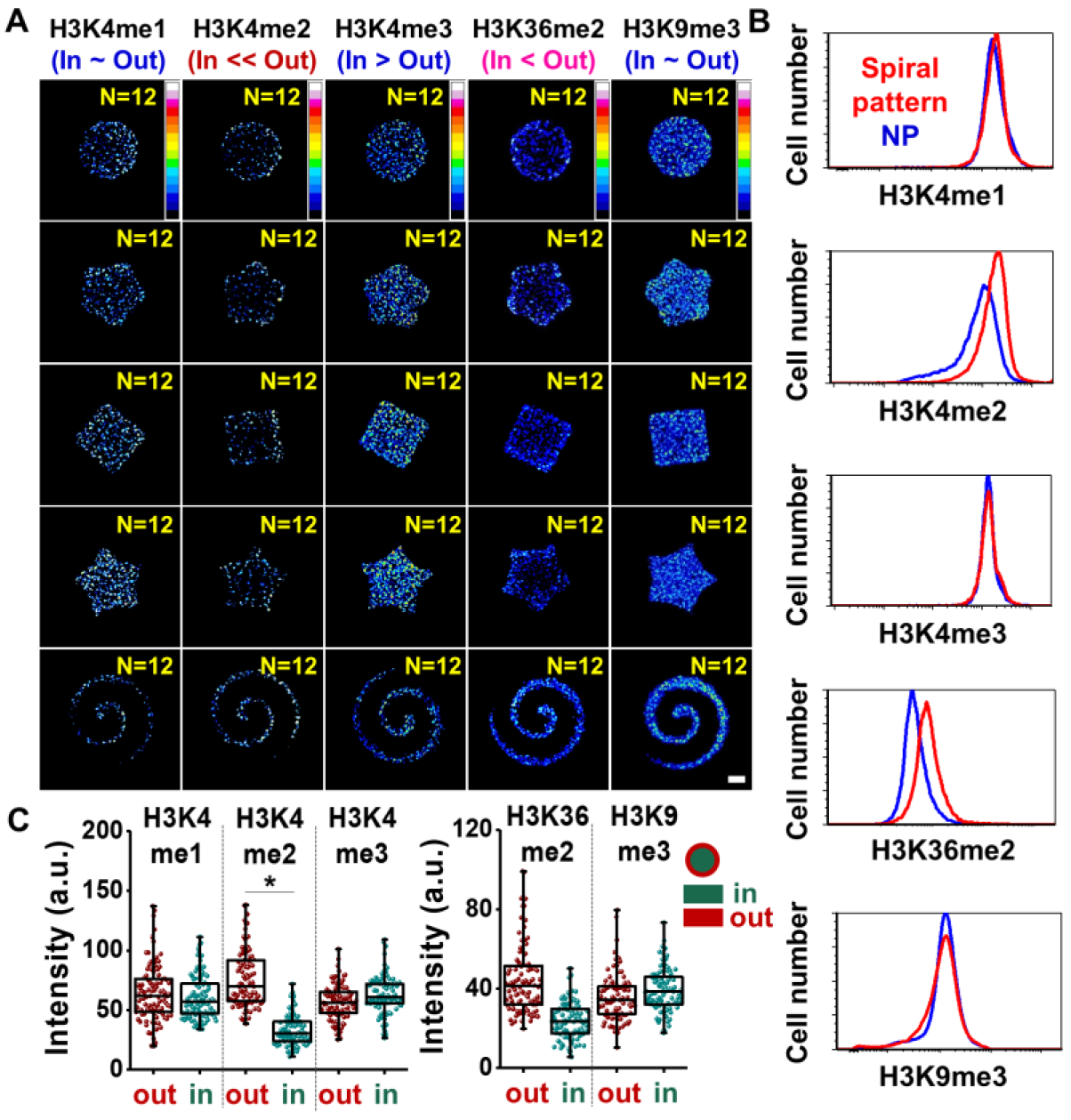
Histone methylation state is influenced by perimeter curvature. (**A**) Immunofluorescence heatmaps of H3K4me3/2/1, H3K36me2, and H3K9me3 for B16F0 cells cultured in a panel of shapes. (**B**) Flow cytometry characterization of histone methylation in B16F0 cells. (**C**) Single cell analysis of histone methylation markers in B16F0 cells cultured in perimeter or central regions of the circular geometry (N=3). Boxes represent 25^th^ to 75^th^ percentile and whiskers represent minimum-maximum. Horizontal lines and points within boxes represent the median and mean respectively for three duplicates. Scale bars, 50 μm. \**P* < 0.05, ANOVA. Error bars represent s.d.

Histone acetylation is also an important modification in regulating chromatin accessibility and regulation of gene expression,^[20]^ and various histone acetylation states have been shown to control gene expression during cancer progression.^[21]^ To probe acetylation activity in our micropatterned cultures, we selected a panel of class I histone deacetylases (HDAC) and measured global acetylation of lysine (AcK), and histone H3 lysine 4 and 9 (H3K4ac and H3K9ac) marks which are associated with gene activation. By applying the same process for identifying methylation states involved in perimeter activation of MICs, we found that cells cultured at the periphery of different shapes expressed higher levels of HDAC1, AcK, H3K4ac, and H3K9ac compared to those cultured at central regions (Fig. 2A, S2, and S4). Flow cytometry of cells cultured in spiral patterns or on non-patterned substrates supported these immunofluorescence results (Fig. 2B). Regional analysis reveals that cells cultured at the periphery of shapes exhibit significant elevation of the H3K9ac mark (Fig. 2C and D), corresponding to lower expression levels of MIC and stemness markers (Fig. S5). AcK and histone marks H3K4ac showed perimeter enhancement, although the difference was not statistically significant. We also immunostained cells cultured along straight lines and ring shapes where curvature and perimeter/area ratio can be varied. After five days in culture, we see cells show higher levels of H3K9ac with increased perimeter curvature and P/A, with a modest reduction in HDAC3 expression, although the difference is not statistically significant. While preliminary, this result supports a potential role for HDAC3 in deacetylating H3K9ac (Fig. 2E).

**Figure 2.**
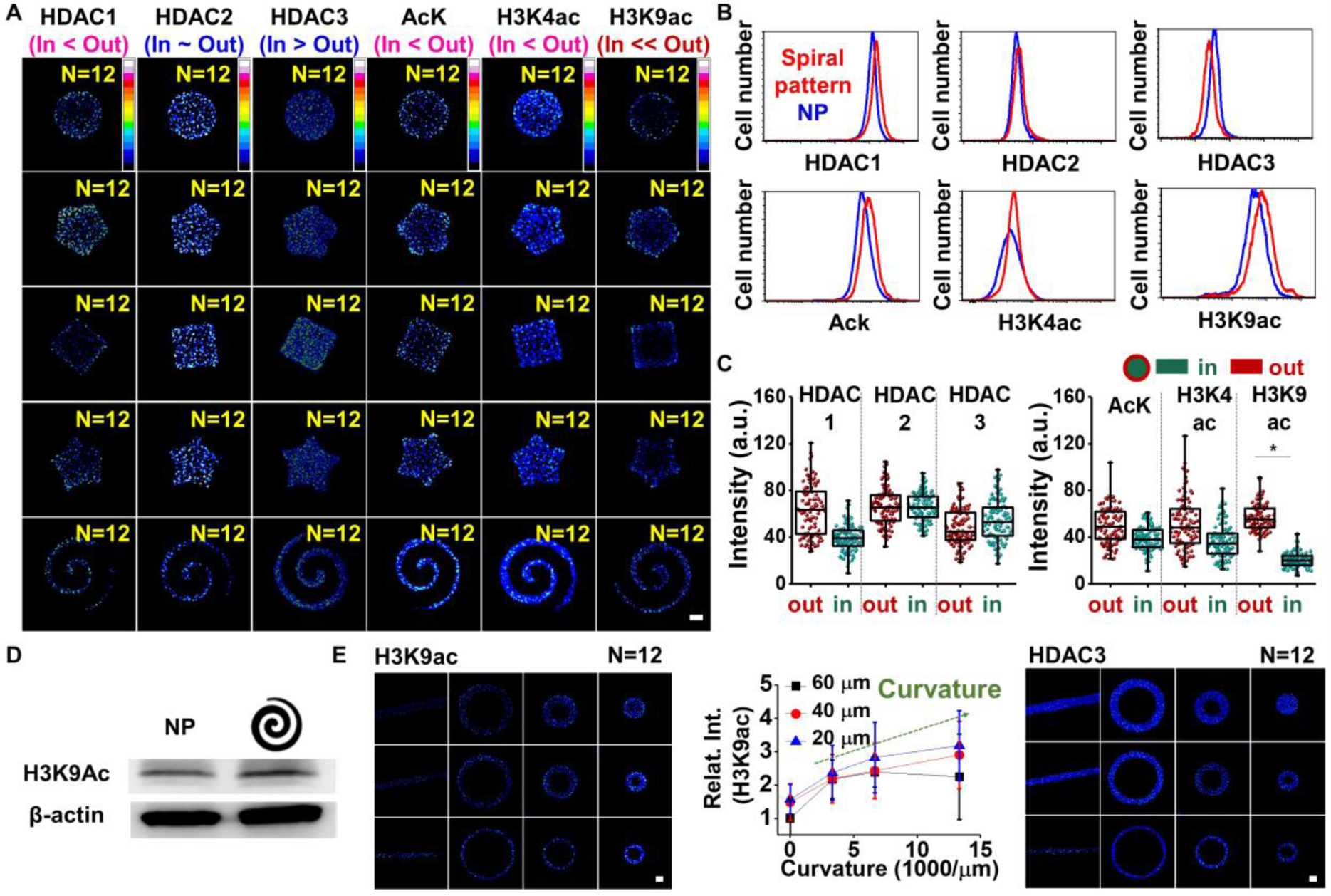
Histone acetylation and deacetylation correspond to epigenetic-mediated phenotype changes in melanoma. (**A**) Immunofluorescence heatmaps of histone acetylation and deacetylation for B16F0 cells cultured on a panel of shapes. (**B**) Flow cytometry characterization of histone acetylation and deacetylation in B16F0 cells. (**C**) Single cell analysis of histone acetylation and deacetylation in B16F0 cells cultured in two different regions of the circular shape (N=3). Boxes represent 25^th^ to 75^th^ percentile and whiskers represent minimum-maximum. Horizontal lines and points within boxes represent the median and mean respectively for three duplicates. Immunofluorescence analysis of histone marks, MIC markers, and transcription factors related to the MIC state over time for cells cultured on (**D**) Western blots for H3K9ac with nonpatterned or spiral-patterned. (**E**) Shapes regulating curvature and perimeter/area to explore the relationship between H3K9ac and HDAC3 (N=3). Scale bars, 50 μm. \**P* < 0.05, ANOVA. Error bars represent s.d.

### 2.2. Inhibiting chromatin-modifying enzymes augment regional variations in the stem cell-like phenotype in melanoma

We next employed an inhibition study in microengineered melanoma aggregates to decouple the potential role of chromatin modifying enzymes in regulating the MIC state. To investigate the role of Jarid1B, a histone demethylase for H3K4me3/2/1 and an established molecular marker for MICs^[15]^ in demethylase activity associated with the stem cell-like state, we cultured B16F0 cells in spiral geometries with small interfering RNA (siRNA) of Jarid1B with scrambled siRNA as control. siRNA concentration and delivery time were adjusted to ensure cells in control and experimental conditions reached approximately the same confluence. After five days in culture, we performed gene expression analysis using quantitative polymerase chain reaction (qPCR) of a panel of markers associated with the MIC state (CD271, Sox2, Oct4, and Nanog). We see a lower degree of transcript expression for markers associated with stemness for cells cultured with Jarid1B siRNA, but we found concentration dependent changes for transcript expression of CD271 (Fig. S6A-F). To evaluate the potential role of Jarid1B in regulating the H3K4me3/2/1 histone marks across spatial regions, we performed immunofluorescence staining of H3K4me3/2/1 for cells cultured in circular shapes, treated with Jarid1B or scrambled siRNA. Jarid1b knockdown does not change the levels of H3K4me2, while leading to an increase in the H3K4me3. This suggests Jarid1b is involved in demethylation of H3K4 but not necessarily associated with regulation of the MIC state at geometric features (Fig. S6G and S7).

Interestingly, we also see a lower degree of transcript expression of HDAC1 for cells cultured with Jarid1B siRNA (Fig. S6E), this may be because HDAC1 is linked to the domains of Jarid1B^[15]^ and one of the EMT-inducing genes (Snail) when complexed with HDAC2.^[22]^ Therefore, to discern a role of HDACs in regulating the MIC phenotype, we supplemented our patterned cultures with the broad spectrum HDAC inhibitors valproic acid (VPA), sodium butyrate (NaB), or Trichostatin A (TSA). Addition of HDAC inhibitors led to a notable increase in not only histone acetylation but also MIC markers (Fig. 3A-C and Fig. S8). The broad-spectrum inhibitory potency of these compounds, coupled with the multivariate roles of HDACs, may give rise to marker dependent variations; however, our hypothesis that histone acetylation augments MIC states remains viable in general and corresponds to a previous report that showed HDAC inhibition played an important role in cancer stem cells and epithelial to mesenchymal transition (EMT).^[23]^ Future work will benefit from the use of more selective inhibitors to isolate the roles of specific enzymes in regulating acetylation state.^[24]^

**Figure 3.**
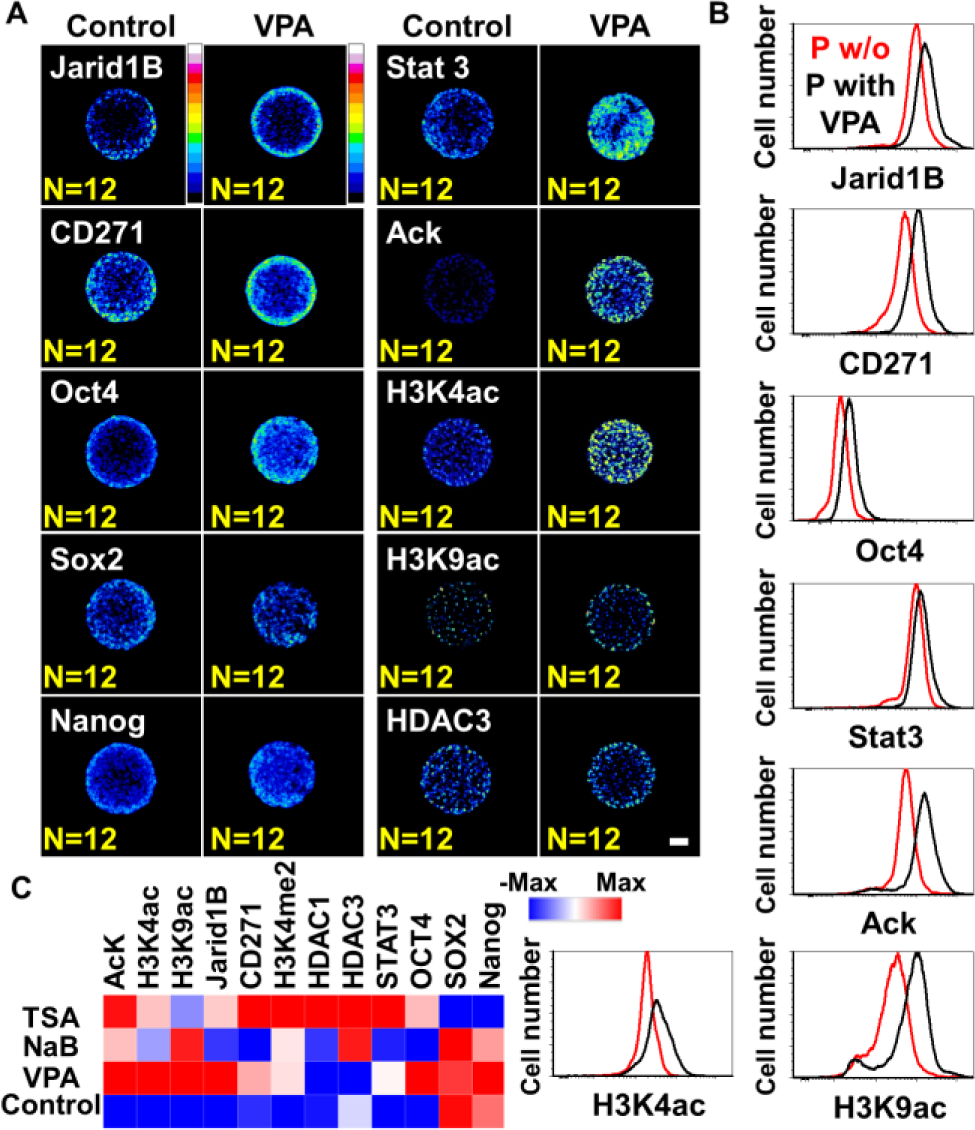
Histone deacetylase activity influences regional variations in the cancer stem cell phenotype in melanoma. (**A**) Immunofluorescence heatmaps of molecular markers associated with the MIC state for B16F0 cells cultured in circular shapes with and without histone deacetylase (HDAC) inhibitors. (**B**) Flow cytometry characterization of these markers. (**C**) Immunofluorescence analysis of molecular markers associated with the MIC state for cells cultured in the presence of multiple HDAC inhibitors (N=3). Scale bars, 50 μm. Error bars represent s.d.

### 2.3. Histone 3 lysine 4 regulates oncogenic gene expression

To understand the possible mechanisms underlying changes in cell state on account of specific histone marks, B16 melanoma cells were grown on spiral patterned (reprogramming) or non-patterned (control) hydrogel substrates for five days, followed by chromatin immunoprecipitation and DNA sequencing (ChIP-seq). ChIP assays specific for the identified histone marks (H3K4me2/H3K9ac) were performed and differential ChIP peaks were identified with at least a 2-fold change and 0.001 FDR cutoff (Fig. 4 and S9). More differential H3K4me2 (57.3%)/H3K9ac (77.8%) peaks were shown for the cells cultured in spiral geometries. To gain insights into potential regulators at these differential sites such as DNA-binding transcription factors, we also performed motif enrichment analysis. We found that differential peaks in spiral patterned versus non-pattered cells are enriched for distinct motif families; *ERG (ETS)*/*Pit1*/*SOX2/9* (reprogramming) or *ETS1*/*TcFap2e1*/*USF2* (control) for H3K4me2 differential peaks and *ERG (ETS)*/*SOX10*/*MITF* (reprogramming) or *RBPJ*/*Nur77*/*Nkx2* (control) for H3K9ac differential peaks. *ETS* genes are known to be linked to p38/ERK mitogen-activated protein kinases (MAPK) signaling for tumor growth and progression.^[25]^ For example, *ETS1* could promote the development and invasion of malignant melanoma,^[26]^ and when it associated with *RhoC* (also enriched for cells on spiral patterned hydrogels), melanoma cells could be progressive and metastatic.^[27]^ Although the ETS family was also a top ranked motif for H3K4me2 peaks in non-patterned cells, enriched gene annotation associated with the differential peaks (Fig. S10 and S11) suggest a distinct role in coordinating the MIC phenotype. *Pit1* is also known to upregulate Snai1, leading to tumor EMT and their growth and metastasis.^[28]^ Similar trends were observed for H3K9ac peaks but it has more distinct and specific differences between cells cultured on spiral patterned and non-patterned hydrogel substrates (Fig. 4).

**Figure 4.**
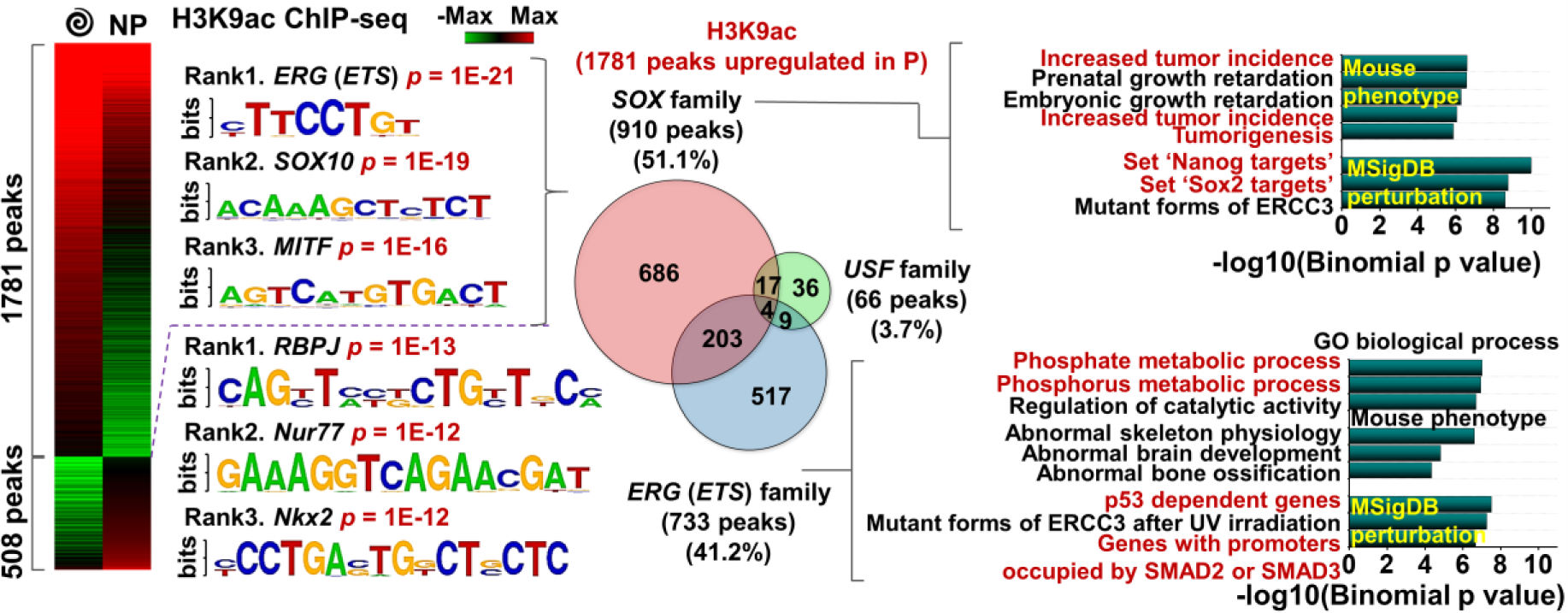
H3K4me2/H3K9ac-regulated gene panels predict phenotypic alterations of melanoma. Heatmap of H3K9ac ChiP-seq signal for cells cultured on spiral geometry or non-patterned substrates. The top three predictive transcription factor motifs with p-values. Enriched annotations of genes for cells cultured on spiral patterns that contain a specific motif (*SOX* or *ETS* family) within the promoter. Venn diagram showing the number of enriched genes for cells cultured on spiral patterns linked to *SOX, ETS*, and *USF* families among H3K9ac-marked genes.

### 2.4. Histone H3 lysine 9 acetylation regulates MIC state through SOX10 motif

One of the top motifs associated with H3K9ac for cells cultured in regions of high curvature and Perimeter/Area is *SOX10*, a neural crest stem cell marker. Previous studies revealed that *SOX10* played an important role in melanoma cell survival, proliferation, and metastasis.^[29]^ It was also reported that the CD271 expression for malanoma, one of representative markers for the MIC state, was directly related to the expression of *SOX10*.^[30]^ In addition, previous studies showed that *MITF* which could function as a melanoma oncogene was associated with melanoma progression,^[31]^ and *SOX10* is known to act upstream of *MITF*,^[32]^ meaning that *SOX10* may thus contribute to the melanoma-specific expression of genes associated with the MIC state. Interestingly, the enriched mouse phenotype annotations related to *SOX10* family in H3K9ac peaks for reprogrammed cells suggest that increased tumor incidence and tumorigenesis are involved in their mouse phenotype. Furthermore, *Nanog* and *SOX2* targets may be perturbed by the *SOX10* family, suggesting the importance of *SOX10* in activation of cells to the MIC state at the tumor periphery. Since we found that the *SOX10* motif was enriched inside the differential histone peaks (Fig. 4 and S12), we conducted immunofluorescence for *SOX10* on our microaggreagates as well as ChIP-seq of *SOX10*. Cells cultured at the perimeter of microaggregates express higher levels of *SOX10* compared to those cultured in central regions (Fig. 5A), and we see 14 differential peaks associated with cells cultured on patterned gels compared to those cultured on non-patterned gels. Note that some genes like Med27 and Trim14 inside H3K4me2 peaks were shown as one of the best differential *SOX10* peaks associated with activated cells, and some peaks located nearby Klf12, Scml4, and Dync1li2 were intersected with differential H3K9 peaks (Fig. S12). These genes may also be involved in malignant melanoma transformation.

**Figure 5.**
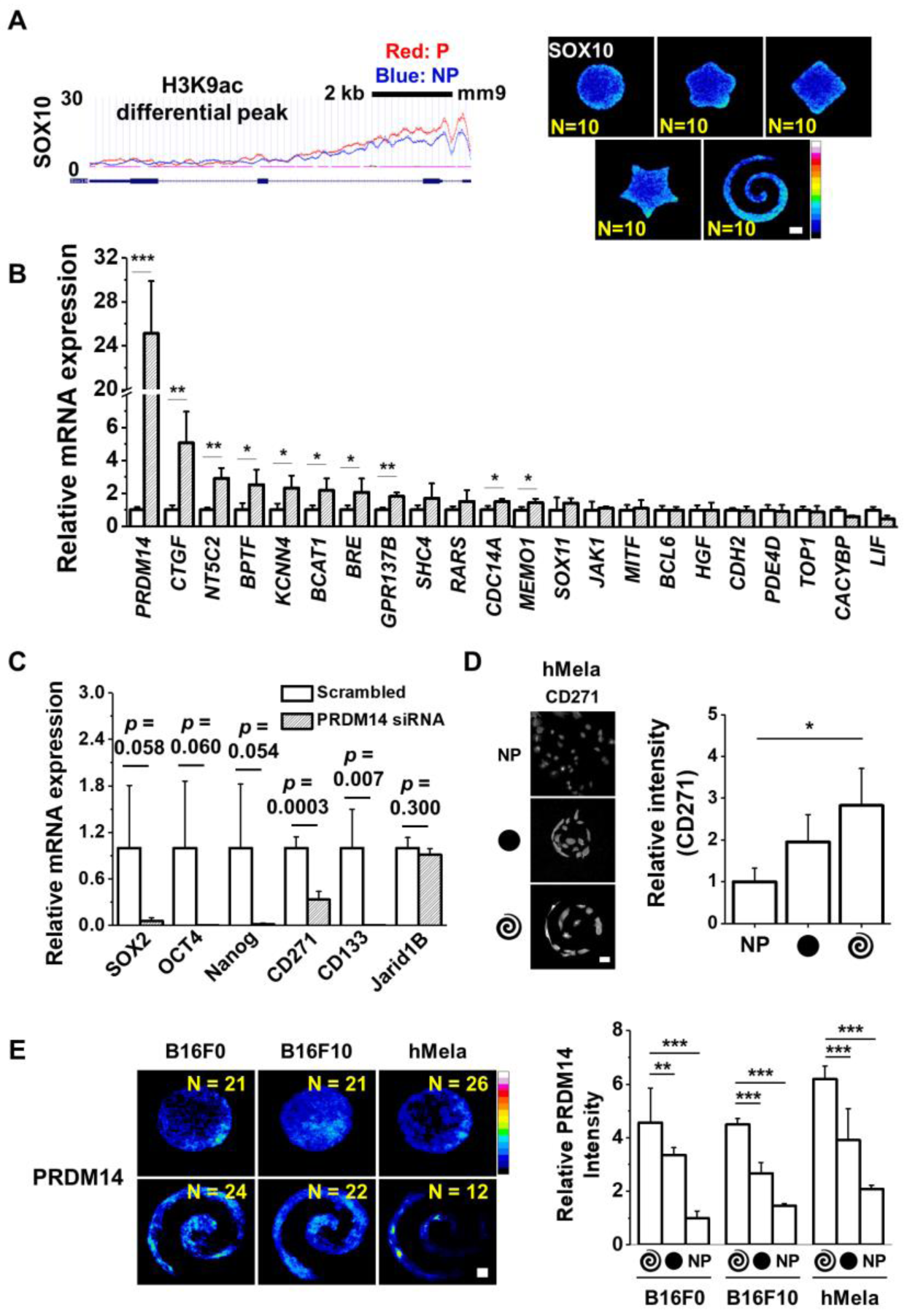
*SOX10* and *PRDM14* are involved in regulating the epigenetic state associated with the MIC phenotype. (**A**) ChiP-seq occupancy for H3K9ac over *SOX10* and immunofluorescence heatmaps of *SOX10* for B16F0 cells cultured in a panel of shapes. (**B**) Results of real-time PCR to measure the expression of genes associated with the differential peaks of H3K9ac/*SOX10* motif. (N=4) (**C**) Results of real-time PCR to measure the gene expression of stemness (*SOX2, OCT4, Nanog*) and MIC (*CD271, CD133, Jarid1B*) for B16F0 cells cultured on spiral geometries for 5 days with PRDM14 or scrambled siRNAs (N=4). (**D**) Representative immunofluorescence images and relative intensity of representative MIC marker, CD271 for hMela cells cultured for five days on circular or spiral geometries or non-patterned substrates. (N=3). (**E**) Immunofluorescence heatmaps and relative intensity of *PRDM14* expression for B16F0, B16F10, and hMela cells cultured for five days on circular or spiral geometries or non-patterned substrates. (N=3). Scale bars, 50 μm. \**P* < 0.05, \*\**P* < 0.005, \*\*\**P* < 0.0005, ANOVA. Error bars represent s.d.

### 2.5. Melanoma initiating cell phenotypes at the perimeter are directed by PR/SET Domain-containing 14 (*PRDM14*)

To further confirm the association between the high ranked regulatory motifs (*ERG (ETS), SOX10*, and *MITF* for activated cells upregulating H3K9ac peaks) and regulation of downstream target genes, we identified H3K9ac differential peaks between two different conditions (reprogrammed and control cells). A number of these differential peaks were located in the regulatory domains of genes associated with cancer growth and progression and thus, we analyzed the expression of these genes for reprogrammed cells over control. Cells were cultured for 5 days on patterned substrates followed by lysis, RNA isolation and real time PCR. Interestingly, activated cells cultured on patterned substrates show significantly higher expression of genes related to malignant melanoma such as CTGF (∼5-fold) and NT5C2 (∼3-fold) compared to cells cultured on non-patterned substrates (Fig. 5B). The most highly differentially expressed gene associated with H3K9ac/SOX10 is *PRDM14* (PR/SET Domain-containing 14, ∼25-fold). *PRDM14* is an epigenetic modifier involved in regulating pluripotency of stem cells with a clear role in modulating expression of core transcription factors Nanog, Sox2, and Oct4.^[33–37]^ *PRDM14* is required to repress genes associated with lineage commitment and ensures naïve pluripotency in embryonic stem cells.^[36]^ In addition, PRDM14 has been implicated in affecting the severity of several human cancers including breast^[38]^ and leukemia^[39]^ with evidence of a link to regulating a stem cell-like state.^[39]^ To date PRDM14 has not been linked to tumorigenicity and the MIC state in melanoma. These findings are supported by the number of overlaps for peak co-occurrence of PRDM14 (pre-existing PRDM14 ChiP-seq data in the literature)^[40]^ with the H3K9ac and H3K4me2 marks from our study within 1000bp, showing significant overlap with all of the peak sets and differential peak sets, except for PvsNP_k4me2 (Table S1).

To explore how *PRDM14* plays a supportive role in coordinating the epigenetic state of melanoma in response to perimeter geometry, we performed knockdowns of *PRDM14* using siRNA. Knockdown was verified by qPCR and Western analysis using three different siRNAs against PRDM14 with a scrambled control (Fig. S13). *PRDM14* knockdown resulted in a decrease in expression for genes linked to the MIC state; knockdown of PRDM14 led to complete abolishment of pluripotency markers *SOX2, OCT4*, and *Nanog*, and putative MIC markers *CD271* and *CD133*; PRDM14 knockdown led to a partial decrease in *Jarid1B* expression (Fig. 5C). To confirm the localization of PRDM14 at perimeter features of micropatterned melanoma aggregates, we performed immunofluorescence characterization in both B16 cells and human primary melanoma cells. Figure 5D shows perimeter enrichment of PRDM14 in the B16F0 cells and to a lesser extent the B16F10 cells. The higher localization at the periphery in the B16F0 cells is consistent with a mechanism where PRDM14 is activated at the interface to reprogram cells of low metastatic potential to a highly metastatic MIC phenotype. To verify our observations in human cells, we chose to look at a BRAF mutant primary human melanoma cell line (hMela). Human melanoma cells show significant enhancement in both CD271 and PRDM14 (Figure 5D and E), suggesting this mechanism is not unique to mouse cell lines but may play a role in guiding the MIC phenotype in human cells. While we have shown how geometric cues can give rise to differential activity of PRDM14, it remains to be demonstrated how PRDM14 is activated based on these biophysical inputs. Furthermore, there are many other histone marks that remain to be profiled in order to map the complexity of epigenetic reprogramming in melanoma. For instance, a recent study demonstrated a link between H3K27me3 and mouse germ cell migration.^[41]^ Nevertheless, our ChIP-seq analysis serves to illuminate how mechanotransduction can regulate the activity of stemness-related epigenetic modifiers including PRDM14 (Fig. 5C).

## 3. Conclusion

In this paper we show how changes in specific histone marks at the perimeter of melanoma aggregates correspond to phenotypic alteration of melanoma cells to a stem cell-like MIC state. We show that stress exerted on microconfined cells at the periphery primes the melanoma phenotype through epigenetic reprogramming via histone modifications H3K9ac and H3K4me2, and involvement of the epigenetic modifier PRDM14. The mechanistic basis of such changes may be related to the response of tumors to their microenvironment where intratumor pressure, extracellular mechanics, and curvature at the margin, coordinate to provide a context in which mechanotransduction and downstream gene expression are regulated by these multivariate signals. Since these microenvironment parameters coincide with tumor angiogenesis,^[19]^ it is important to consider the ramifications of transformation to a MIC state in proximity to pathways for metastasis. These findings may help guide researchers in further exploring epigenetic signatures for tumor malignancy, and the development of novel strategies to prevent, diagnose, and treat metastatic cancers.

## 4. Experimental Section

### Hydrogel fabrication

10 kPa polyacrylamide hydrogels (PA) were made as described previously.^[14]^ Briefly, 10% acrylamide and 0.1% bis-acrylamide (Sigma) solution were prepared and mixed with initiators, 0.01% ammonium persulfate (APS, Sigma) and tetramethylethylenediamine (TEMED, Sigma), to initiate gelation. 20 µl of the mixture was sandwiched between a glass coverslip (18 mm, Fisher Scientific) functionalized with 3-aminopropyltriethoxysilane for 3 min and glutaraldehyde for 30 min (Sigma) and a hydrophobically treated glass slide to generate the even and homogeneous surface of gels on the activated coverslips. After around 25 min of gelation, the coverslips conjugated with gels were gently detached from the hydrophobically treated glass slide. Hydrazine hydrate (55%) chemistry was employed to modify the surface chemistry of PA gels and applied to the gels surface for 2 h with rocking, followed by washing with 5% glacial acetic acid for 1 h and distilled water for 1 h. PA gels fabricated on cover slips were stored at 4°C for later use. Polydimethylsiloxane (PDMS, Polysciences) stamps were fabricated from silicon masters made by conventional photolithography for patterned or non-patterned shapes. To generate free aldehydes from oxidize sugar groups in matrix protein (fibronectin, Sigma), sodium periodate (∼3.5mg/ml, Sigma) for at least 45 min was employed. The protein solution was mounted onto patterned or non-patterned (flat surface) stamps for 30 min and dried with air. Micro-contact printing was used to transfer the protein residues on stamps to the gel surface (chemical conjugation).

### Cell source and culture

The cancer cell lines B16F0 cells were purchased from American Type Culture Collection (ATCC) and cultured according to the recommended protocols. Media was changed every 3 to 4 days and cells were passaged at nearly 90% confluence using 0.25% trypsin (Gibco). B16F0 cells were verified for mycoplasma contamination at Charles River Laboratories for cell line testing. De-identified primary human melanoma cells were a kind gift from John A. Copland III from the Mayo Clinic, Jacksonville, FL.

### Immunofluorescence

B16F0 cells were fixed using 4% paraformaldehyde for 20 minutes. Cells were permeabilized with 0.1% Triton-X100 for 30 min at room temperature and then blocked with 1% bovine serum for 15 min. Cells were stained with the appropriate primary and secondary antibodies (Table S2). Before every step, cells were washed at least twice with PBS. Imaging was performed using an LSM 700 (Carl Zeiss, Inc.) four laser point scanning confocal microscope with a single pinhole for confocal imaging for fluorescence imaging.

### RNA isolation and RT-PCR

Adherent cells on patterned gels (12 identical substrates for each condition) were lysed directly in TRIZOL reagent (Invitrogen). Total RNA was isolated by chloroform extraction and ethanol precipitation and amplified using TargetAmp™ 1-Round aRNA Amplification Kit 103 (Epicentre) according to vendor protocols. Superscript III^®^ First Strand Synthesis System for RT-PCR (Invitrogen) was employed to reversely transcribe total RNA. RT-PCR was performed using SYBR^®^ Green Real-Time PCR Master Mix (Invitrogen) on an Eppendorf Realplex 4S Real-time PCR system. Primer sequences were in supplementary Table 3. All reactions were performed linearly by cycle number for each set of primers.

### Western analysis

Cell extracts were isolated using RIPA buffer supplemented with protease and phosphatase inhibitors. Protein concentrations were determined by Nanodrop or BCA protein assay (Thermofisher), according to the company instructions. Subsequently, proteins were separated by 4-20% SDS-PAGE and electrophoretically transferred onto PVDF (Thermo Fisher Scientific) or nitrocellulose membranes (Bio-rad, Australia), which were then probed with primary antibodies overnight at 4 °C. HRP-conjugated secondary antibodies were detected by chemiluminescence agent ECL or Supersignal Western Dura Extended Duration (Thermofisher, Australia). Membranes were imaged by ImageQuant LAS4000 (GE healthcare, Sweden). Densitometric analysis was performed by ImageQuant TL Software (GE healthcare) and presented as ratios of protein expression normalized to relevant GAPDH or β-actin loading control.

### Inhibition assay and siRNA

Histone deacetylase (HDAC) inhibitors, valproic acid (VPA, Sigma-Aldrich (P4543)) sodium butyrate (NaB, Bio Vision (1609-1000)), or Trichostatin A (TSA, Sigma-Aldrich (T8552)) were added to cells in media before seeding and after changing media at 1 μg/ml, respectively. MAP kinase inhibitors for ERK1/2 (FR180204) and p38 (SB202190) (Calbiochem) were supplemented in the media at 6 μM after seeding cells and changing each media. Blocking integrin α5ß1 was performed by adding the antibodies to cells in media before seeding at 1μg/ml. The siRNAs for Jarid1B (ID 75605, Trilencer-27 Mouse siRNA, siRNA A: SR422988A, siRNA B: SR422988B, and siRNA C: SR422988C) or scrambled siRNAs (SR30004) were purchased from OriGene. Transfection was performed according to the vendor’s instructions. Lipofectamine 2000™was employed for higher transfection efficiency. Cells cultured for 5 days in patterned substrates were treated with siRNA twice at day 1 and day 3.

### Cell labelling and flow cytometry

B16F0 cells cultured for five days on spiral-patterned or non-patterned gels (12 identical substrates for each condition) were isolated from substrates by trypsin, followed by breaking down into a single cell suspension. Cells were fixed in 4% paraformaldehyde for 20 min and then permeabilized in 0.1% Triton X-100 in PBS for 30 min. After blocking cells in 1% BSA in PBS for 1 h, Cells were stained with primary antibodies in 1% BSA in PBS overnight at 4°C and then secondary antibodies in 2% goat serum, 1% BSA in PBS for 20 min in a humid chamber (5% CO2 and 37°C). Before every step, cells were rinsed at least three times with PBS. A BD LSR Fortessa Flow Cytometry Analyzer was used to perform flow cytometry analysis. To set the baseline, negative controls were prepared by staining cells without primary antibodies.

### Microscopy data analysis

Confocal images were analyzed using ImageJ software. Multiple cells were imaged for each condition and fluorescence intensities of single cells in different regions of patterns (after background subtraction) were used to compare marker expression. For generating immunofluorescence heatmaps, cells cultured on various shapes were fixed, stained, and imaged on the same day using the same settings. After subtraction of background intensities of raw fluorescent images, patterns were aligned in ImageJ with the same orientation as cultured across the surface, followed by incorporating into a Z stack with the average intensity calculated for heatmap generation.

### Chromatin immunoprecipitation and sequencing (Chip-seq)

H3K4me2 and H3K9ac ChiP samples were prepared from B16 melanoma cells cultured on patterned or non-patterned substrates, and ChiP DNA quality was verified as previously described^41^. B16 melanoma cells were cultured for five days and then fixed with 1% formaldehyde for 10 min at room temperature. Fixations were quenched by glycine (125 mM), followed by washing cells with cold 1x PBS two times. Cells were treated with hypotonic lysis buffer for 10 min (20 Mm HEPES at pH 7.9, 10 mM KCl, 1 mM EDTA at pH 8, 10% glycerol, 1 mM DTT, 0.5 mM PMSF, 0.1 mM sodium orthovanadate, and 1X Roche protease inhibitors). Collected nuclear pellets were lysed in in 1× RIPA buffer (10 mM Tris-Cl at pH 8.0, 140 mM NaCl, 1% Triton X-100, 0.1% SDS, 1% deoxycholic acid, 0.5 mM PMSF, 1 mM DTT, 0.1 mM sodium orthovanadate, and Roche protease inhibitors). Nuclear lysates were sonicated with a Branson 250 Sonifier (output 20%, 100% duty cycle) to shear the chromatin to ∼1 Kb in size. Clarified lysates were incubated overnight at 4°C with anti-H3K4me2 (Cell Signaling, 9725) or H3K9ac (Cell Signaling, 9649) antibodies. Protein–DNA complexes were precipitated, immunoprecipitates were washed three times in 1× RIPA, once in 1× PBS, and then eluted from the beads by addition of 1% SDS, 1× TE (10 mM Tris-Cl at pH 7.6, 1 mM EDTA at pH 8), and incubated for 10 min at 65°C. Cross-links were reversed overnight at 65°C. Purification for all samples were performed by treatment first with 200 μg/mL RNase A for 1 h at 37°C, then with 200 μg/mL Proteinase K for 2 h at 45°C, followed by extraction with phenol:chloroform:isoamyl alcohol and precipitation at -70°C with 0.1 volume of 3 M sodium acetate, 2 volumes of 100% ethanol, and 1.5 μL of pellet paint coprecipitant. ChIP DNA prepared from 1 × 107 cells was resuspended in 50 μL of ultrapure water. Sequencing (an Illumina HiSeq 2500 sequencer using a TruSeq SBS sequencing kit, version 4) was performed and Fastq files were obtained and demultiplexed with the bcl2fastq v2.17.1.14 Conversion Software (Illumina). The sequencing data from this study have been submitted to the NCBI Gene Expression Omnibus (GEO; http://www.ncbi.nlm.nih.gov/geo/).

### Chip-seq data analysis

ChIP-seq bioinformatics analyses were performed as previously described^41^. Sequence data were mapped with Bowtie2 (Langmead and Salzberg 2012) to the UCSC Mus musculus mm9 genome, using default settings. Mapped sequence data were analyzed for peaks using HOMER (Hypergeometric Optimization of Motif EnRichment) v4.7 (Heinz et al. 2010). Samples were converted into tag directories, and QC was performed using read mapping and GC bias statistics. Histone peaks were then called from the Tag Directories with default factor settings, except local filtering was disabled (-L 0) and input filtering was set at three-fold over background (-F 3), to increase the sensitivity of the peak calling and identify individual subunits of multihistone peaks. After peak calling, peak files were annotated to the mouse mm9 genome using HOMER’s annotation script to assign peaks to genes, and associate peaks with differential expression data. BigWiggle pileup files were generated using HOMER’s makeBigWig.pl script with default settings. Differential chromatin peaks were identified using the HOMER getDifferentialPeak.pl script, looking for any peaks that changed at least two-fold between conditions with a significance cutoff of 1 × 10^−4^. Genes annotated nearby differential peaks were submitted for GO analysis to DAVID and GREAT (Dennis et al. 2003; McLean et al. 2010). Motif Analysis was performed with the HOMER findMotifsGenome.pl script using default settings.

### Statistical analysis

Data were obtained at least three independent experiments and error bars represent standard deviation around the mean unless otherwise specified. For comparing statistics between two groups or more than two groups, student’s *t*-test or analysis of variance (ANOVA) with Tukey HSD Post-hoc testing, respectively, were employed. Differences were considered significant at *P*<0.05.

## Acknowledgements

This research was supported with funding from the American Cancer Society Illinois Division Grant # 281225, the National Science Foundation Grant # 1454616 CAR, and the Australian Research Council Grant # FT180100417. CHS was supported by #SFLife 291812 from the Simons Foundation. We gratefully acknowledge the support by Prof. Lisa J. Stubbs from the Department of Cell and Developmental Biology, University of Illinois at Urbana-Champaign. We also thank the Institute of Genomic Biology Imaging facilities, Micro, Nanotechnology Laboratory facilities, and the Roy J. Carver Biotechnology Center.

## References

[1] P. C. Nowell, Science 1976, 194, 23.

[2] M. Esteller, Adv. Exp. Med. Biol. 2003, 532, 39.

[3] A. D. Goldberg, C. D. Allis, E. Bernstein, Cell 2007, 128, 635.

[4] A. P. Feinberg, B. Tycko, Nat. Rev. Cancer 2004, 4, 143.

[5] J. T. L. Leightone J. Core, Josua J. Waterfall, Science 2008, 322, 1845.

[6] B. Schuettengruber, D. Chourrout, M. Vervoort, B. Leblanc, G. Cavalli, Cell 2007, 128, 735.

[7] T. Schatton, G. F. Murphy, N. Y. Frank, K. Yamaura, A. M. Waaga-Gasser, M. Gasser, Q. Zhan, S. Jordan, L. M. Duncan, C. Weishaupt, R. C. Fuhlbrigge, T. S. Kupper, M. H. Sayegh, M. H. Frank, Nature 2008, 451, 345.

[8] J. E. Visvader, G. J. Lindeman, Nat. Rev. Cancer 2008, 8, 755.

[9] C. E. Meacham, S. J. Morrison, Nature 2013, 501, 328.

[10] A. B. Hjelmeland, Q. Wu, J. M. Heddleston, G. S. Choudhary, J. MacSwords, J. D. Lathia, R. McLendon, D. Lindner, A. Sloan, J. N. Rich, Cell Death Differ. 2011, 18, 829.

[11] C. Lagadec, E. Vlashi, L. Della Donna, C. Dekmezian, F. Pajonk, Stem Cells 2012, 30, 833.

[12] M. F. Pang, M. J. Siedlik, S. Han, M. Stallings-Mann, D. C. Radisky, C. M. Nelson, Cancer Res. 2016, 76, 5277.

[13] J. M. Heddleston, Z. Li, R. E. McLendon, A. B. Hjelmeland, J. N. Rich, Cell Cycle 2009, 8, 3274.

[14] J. Lee, A. A. Abdeen, K. L. Wycislo, T. M. Fan, K. A. Kilian, Nat. Mater. 2016, 15, 856

[15] B. J. Klein, L. Piao, Y. Xi, H. Rincon-arano, B. Scott, D. Peng, H. Wen, C. Larson, X. Zhang, X. Zheng, A. Michael, P. V Peña, A. Mangan, D. L. Bentley, B. D. Strahl, Cell Rep. 2015, 6, 325.

[16] Q. Li, L. Shi, B. Gui, W. Yu, J. Wang, D. Zhang, X. Han, Z. Yao, Y. Shang, Cancer Res. 2011, 71, 6899.

[17] A. Roesch, M. Fukunaga-Kalabis, E. C. Schmidt, S. E. Zabierowski, P. A. Brafford, A. Vultur, D. Basu, P. Gimotty, T. Vogt, M. Herlyn, Cell 2010, 141, 583.

[18] M. Yoshida, A. Ishimura, M. Terashima, Z. Enkhbaatar, N. Nozaki, K. Satou, T. Suzuki, Biochem. J. 2011, 437, 555.

[19] J. Lee, A. A. Abdeen, J. Hedhli, K. L. Wycislo, I. T. Dobrucki, T. M. Fan, L. W. Dobrucki, K. A. Kilian, Sci. Adv. 2017, 3, e1701350.

[20] M. D. Shahbazian, M. Grunstein, Annu. Rev. Biochem. 2007, 76, 75.

[21] S. Sharma, T. K. Kelly, P. A. Jones, Carcinogenesis 2009, 31, 27.

[22] J. von Burstin, S. Eser, M. C. Paul, B. Seidler, M. Brandl, M. Messer, A. von Werder, A. Schmidt, J. Mages, P. Pagel, A. Schnieke, R. M. Schmid, G. Schneider, D. Saur, Gastroenterology 2009, 137, 361.

[23] F. S. Giudice, D. S. Pinto, J. E. Nor, C. H. Squarize, R. M. Castilho, PLoS One 2013, 8, e58672.

[24] J. Roche, P. Bertrand, Eur. J. Med. Chem. 2016, 121, 451.

[25] P. C. Hollenhorst, M. W. Ferris, M. A. Hull, H. Chae, S. Kim, B. J. Graves, Genes Dev. 2011, 25, 2147.

[26] T. Rothhammer, J. C. Hahne, A. Florin, I. Poser, F. Soncin, N. Wernert, A. K. Bosserhoff, Cell Mol Life Sci 2004, 61, 118.

[27] B. Spangler, M. Kappelmann, B. Schittek, S. Meierjohann, L. Vardimon, A. K. Bosserhoff, S. Kuphal, Int. J. Cancer 2012, 130, 2801.

[28] I. Ben-batalla, S. Seoane, T. Garcia-caballero, R. Gallego, M. Macia, L. O. Gonzalez, F. Vizoso, R. Perez-fernandez, J. Clin. Invest. 2010, 120, 4289.

[29] K. B. Tudrej, E. Czepielewska, M. Kozłowska-wojciechowska, Arch. Med. Sci. 2016, 133, 1493.

[30] O. Shakhova, D. Zingg, S. M. Schaefer, L. Hari, G. Civenni, J. Blunschi, S. Claudinot, M. Okoniewski, F. Beermann, D. Mihic-Probst, H. Moch, M. Wegner, R. Dummer, Y. Barrandon, P. Cinelli, L. Sommer, Nat. Cell Biol. 2012, 14, 882.

[31] L. A. Garraway, H. R. Widlund, M. A. Rubin, G. Getz, A. J. Berger, S. Ramaswamy, R. Beroukhim, D. A. Milner, S. R. Granter, J. Du, C. Lee, S. N. Wagner, C. Li, T. R. Golub, D. L. Rimm, M. L. Meyerson, D. E. Fisher, W. R. Sellers, Nature 2005, 436, 117.

[32] S. B. Potterf, M. Furumura, K. J. Dunn, H. Arnheiter, W. J. Pavan, Hum. Genet. 2000, 107, 1.

[33] A. Burton, J. Muller, S. Tu, P. Padilla-Longoria, E. Guccione, M. E. Torres-Padilla, Cell Rep. 2013, 5, 687.

[34] Z. Ma, T. Swigut, A. Valouev, A. Rada-Iglesias, J. Wysocka, Nat. Struct. Mol. Biol. 2011, 18, 120.

[35] A. Gillich, S. Bao, N. Grabole, K. Hayashi, M. W. B. Trotter, V. Pasque, E. Magnúsdóttir, M. A. Surani, Cell Stem Cell 2012, 10, 425.

[36] M. Yamaji, Y. Seki, K. Kurimoto, Y. Yabuta, M. Yuasa, M. Shigeta, K. Yamanaka, Y. Ohinata, M. Saitou, Nat. Genet. 2008, 40, 1016.

[37] N. Okashita, Y. Suwa, O. Nishimura, N. Sakashita, M. Kadota, G. Nagamatsu, M. Kawaguchi, H. Kashida, A. Nakajima, M. Tachibana, Y. Seki, Stem Cell Reports 2016, 7, 1072.

[38] N. Nishikawa, M. Toyota, H. Suzuki, T. Honma, T. Fujikane, T. Ohmura, T. Nishidate, M. Ohe-Toyota, R. Maruyama, T. Sonoda, Y. Sasaki, T. Urano, K. Imai, K. Hirata, T. Tokino, Cancer Res. 2007, 67, 9649.

[39] E. J. Dettman, S. J. Simko, B. Ayanga, B. L. Carofino, J. F. Margolin, H. C. Morse, M. J. Justice, Oncogene 2011, 30, 2859.

[40] N. Y. Chia, Y. S. Chan, B. Feng, X. Lu, Y. L. Orlov, D. Moreau, P. Kumar, L. Yang, J. Jiang, M. S. Lau, M. Huss, B. S. Soh, P. Kraus, P. Li, T. Lufkin, B. Lim, N. D. Clarke, F. Bard, H. H. Ng, Nature 2010, 468, 316.

[41] A. Mallol, M. Guirola, B. Payer, Epigenetics Chromatin. 2019, 12, 38.

