## Supplemental Figure S1-S13 and Table S1-S4 for "Geometric regulation of histone state directs melanoma reprogramming"

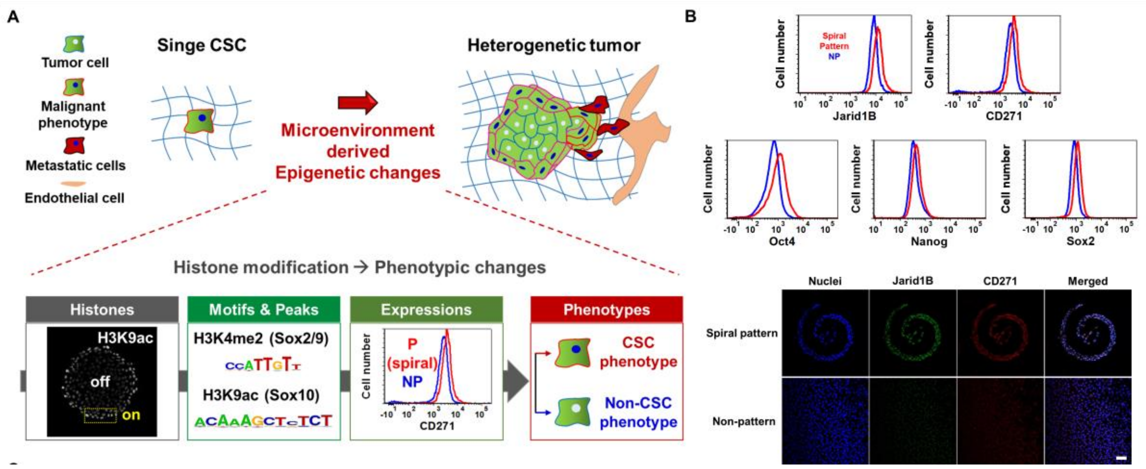


**Fig. S1.** (**A**) Scheme of phenotypic alterations caused by histone modifications in response to microenvironment derived epigenetic changes. (**B**) Flow cytometry characterization of MIC (Jarid1B and CD271) and stemness (Oct4, Nanog, and Sox2) markers in B16F0 cells cultured on spiral patterned or non-patterned substrates. Representative confocal images of Jarid1B and CD271 for B16F0 cells cultured on spiral patterned or non-patterned substrates.

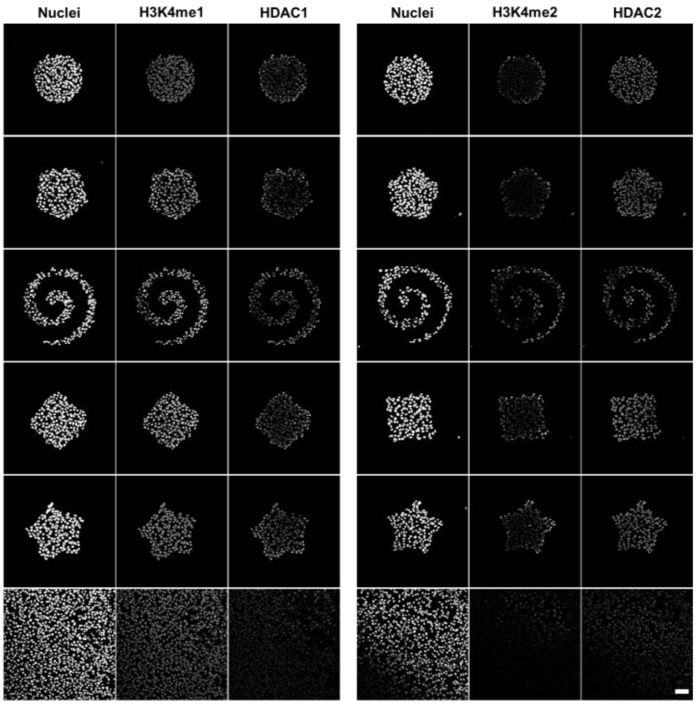
Nuclei
H3K4me1
HDAC1
Nuclei
H3K4me2
HDAC2

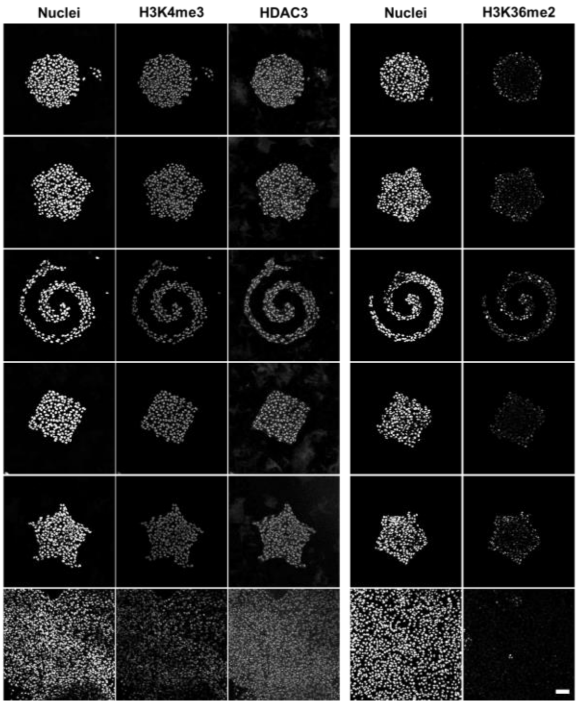
Nuclei
H3K4me3
HDAC3
Nuclei
H3K36me2

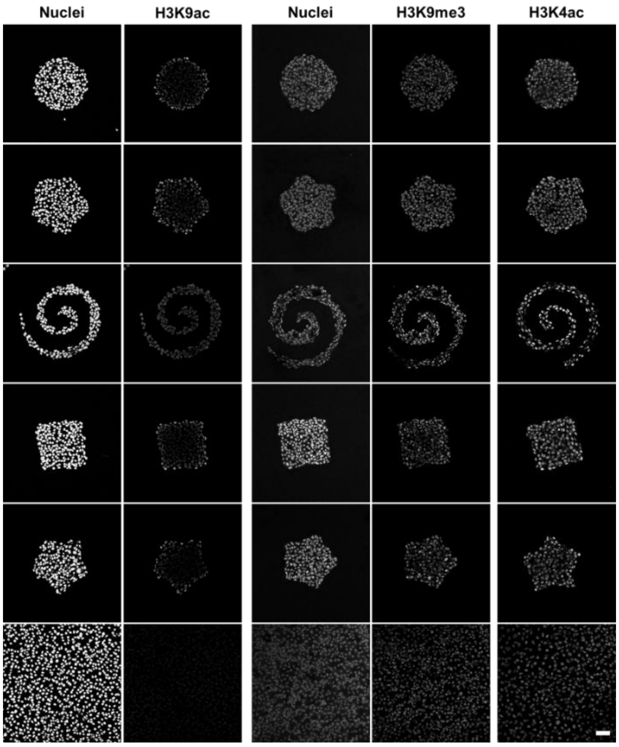


**Fig. S2.** Representative confocal images of methylation (H3K4me3/2/1, H3K9me3, and H3K36me2), HDAC1/2/3, and acetylation (H3K4ac and H3K9ac) for B16F0 cells cultured in a panel of shapes.

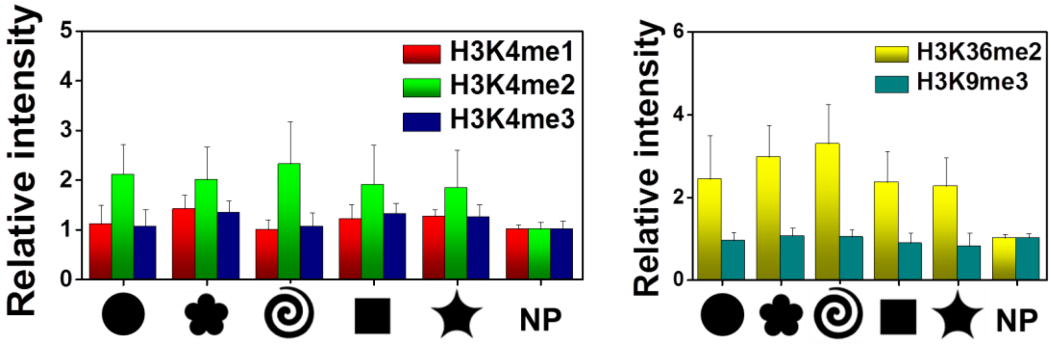


**Fig. S3.** Histone expression of methylation markers for B16F0 cells cultured in a panel of shapes or non-patterned substrates (N=3). Error bars represent s.d.

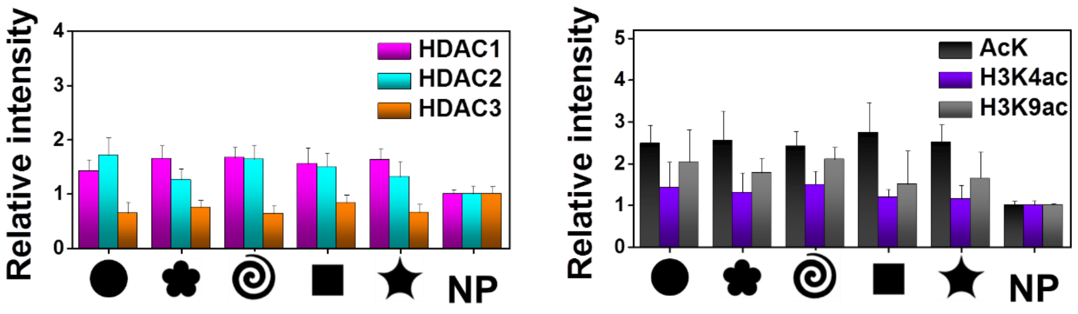


**Fig. S4.** Intensity of acetylation markers for B16F0 cells cultured in a panel of shapes or non-patterned substrates (N=3). Error bars represent s.d.

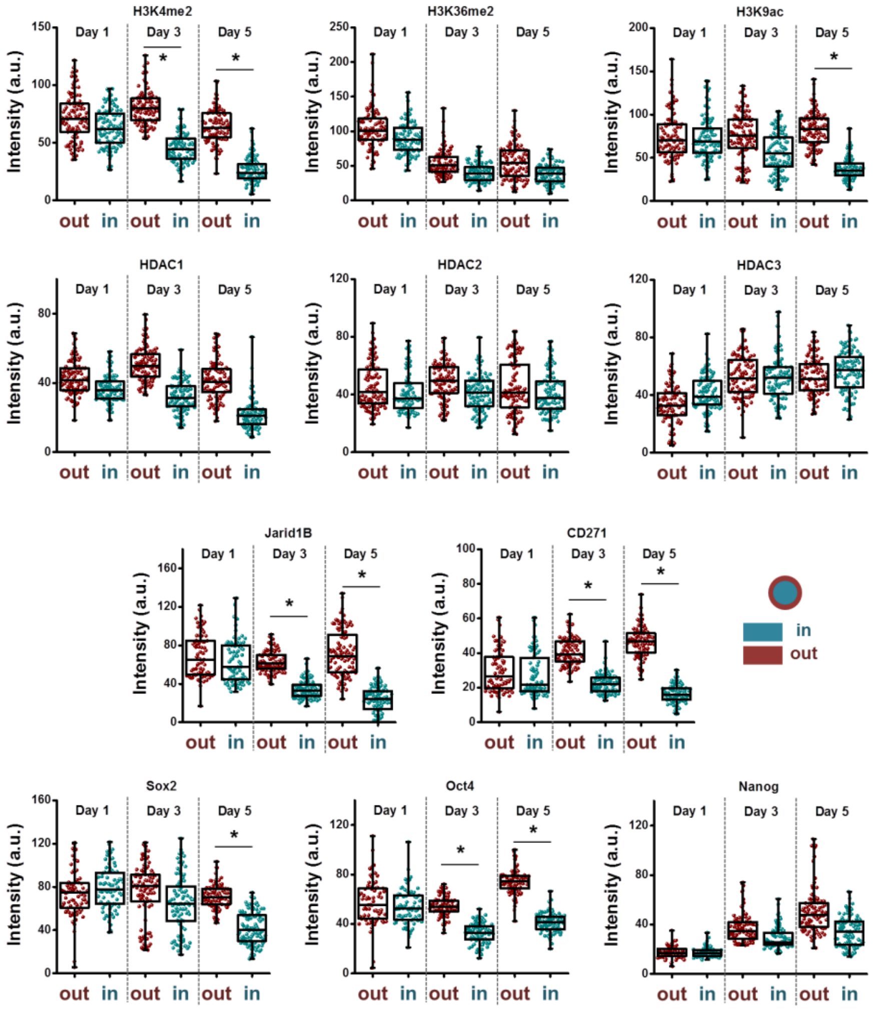


**Fig. S5.** Intensity of Histone modifications (H3K4me2, H3K36me2, H3K9ac, and HDAC1/2/3), MIC markers (Jarid1B and CD271), and transcriptional factors related to stemness and MIC state (Sox2, Oct4, and Nanog) depending on culture time (day 1, 3, and 5) for cells cultured on different regions (outside/inside ratio) of circular shape (N=3). Error bars represent s.d.

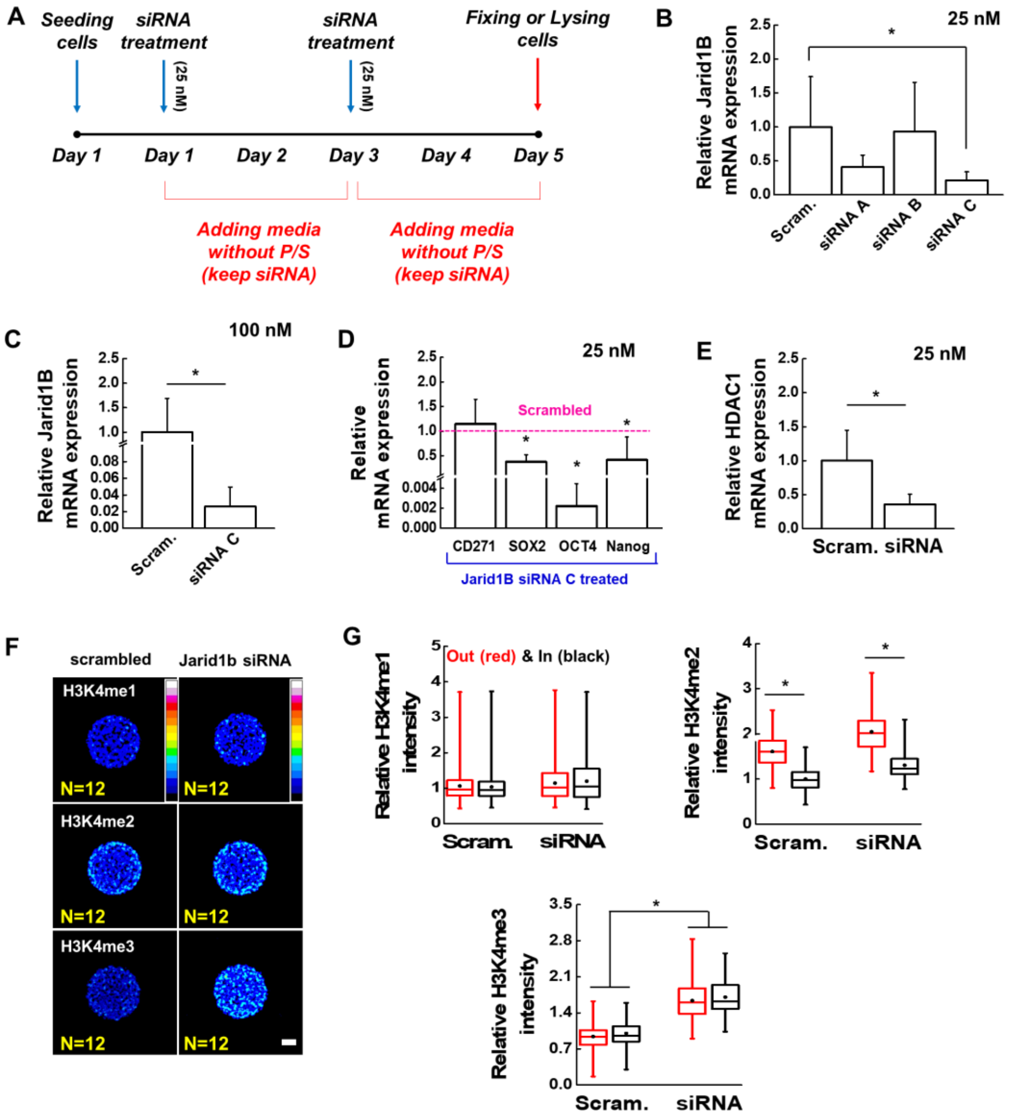


**Fig. S6.** (**A**) Media conditions for normal, inhibition, or siRNA transfection. Results of real-time PCR to measure the gene expression of Jarid1B (three different sequences of Jarid1B siRNAs (A, B, and C) with different concentrations, (**B**) 25 or (**C**) 100 nM. (**D**) Results of real-time PCR to measure the gene expression of CD271, Sox2, Oct4, and Nanog for B16F0 cells cultured on spiral geometries for 5 days with Jarid1B or scrambled siRNAs and (**E**) HDAC1 for cells cultured on spiral geometry for 5 days with Jarid1B or scrambled siRNAs (N=5). (**F**) Immunofluorescence heatmaps of H3K4me3/2/1expression for B16F0 cells cultured on circular geometries treated with scrambled or Jarid1B siRNA. (**G**) Jarid1B regulates the levels of demethylation of H3K4me3/2/1 with different efficiencies (N=3). Error bars represent s.d.

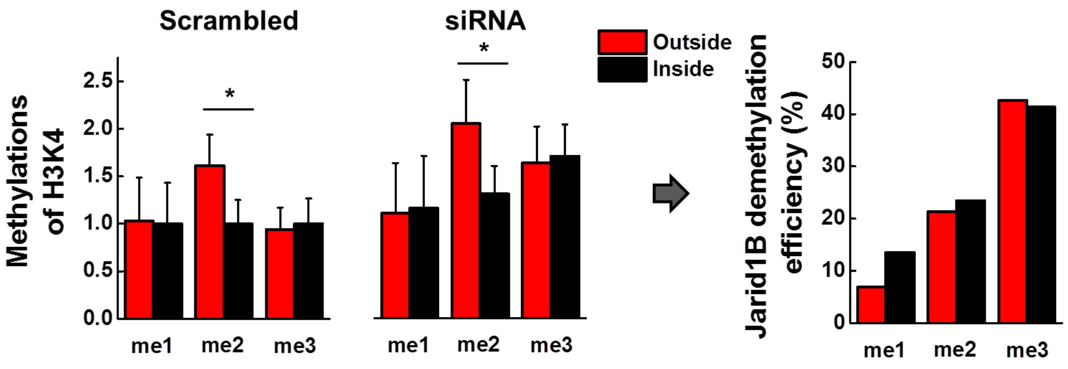


**Fig. S7.** Histone H3K4me3/2/1 expression for B16F0 cells cultured in perimeter or central regions of the circular geometry (N=3), and calculated Jarid1B demethylation efficiency through the composition between cells cultured with scrambled and Jarid1B siRNA. Error bars represent s.d.

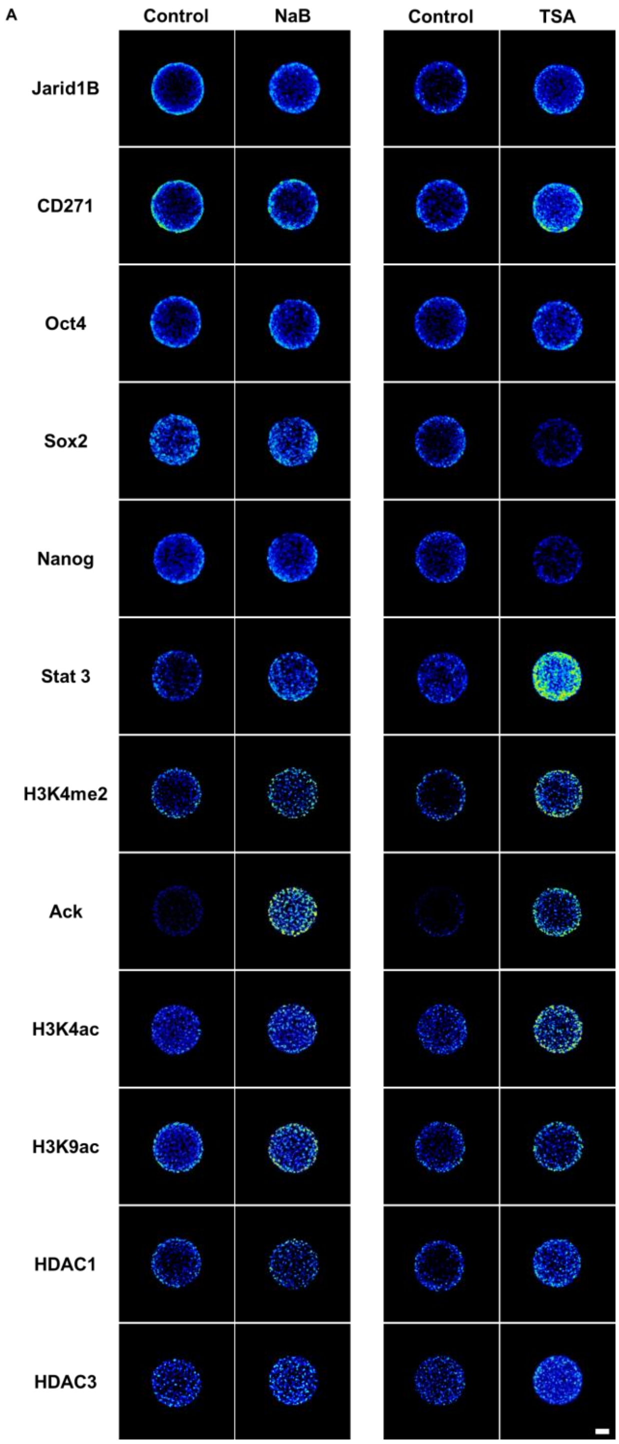
A
Control
NaB
Control
TSA
Jarid1B
CD271
Oct4
Sox2
Nanog
Stat 3
H3K4me2
Ack
H3K4ac
H3K9ac
HDAC1
HDAC3

B

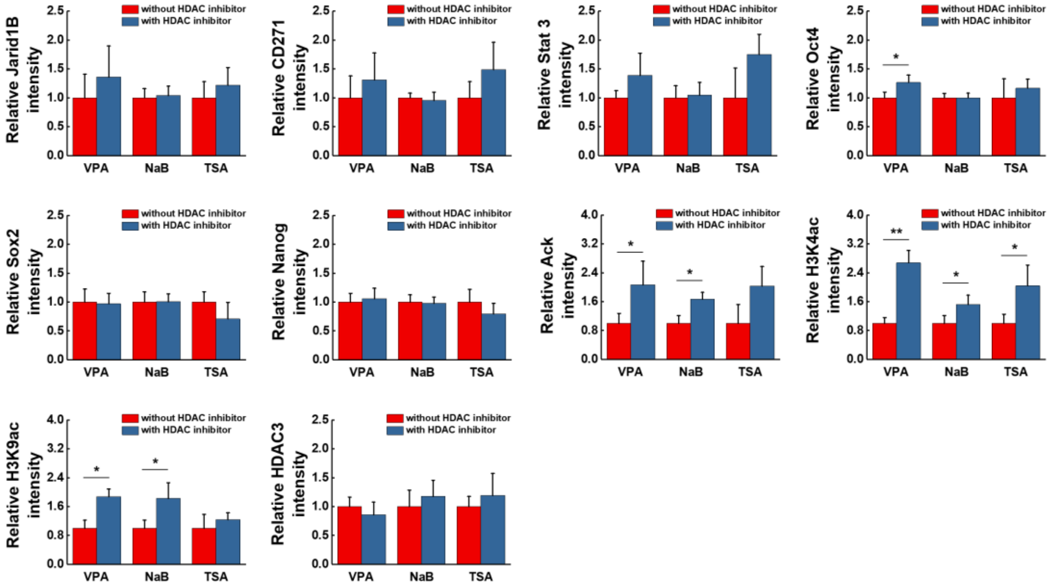


**Fig. S8.** (**A**) Immunofluorescence heatmaps of histone acetylation and deacetylation, CSC surface marker (CD271), transcriptional factors related to stemness and CSC state for B16F0 cells cultured in circular shapes (N=12). (B) Relative immunofluorescence intensity of the markers we selected for cells cultured in spiral geometry with/without HDAC inhibitors (N=3). Scale bars, 50 μm. Error bars represent s.d.

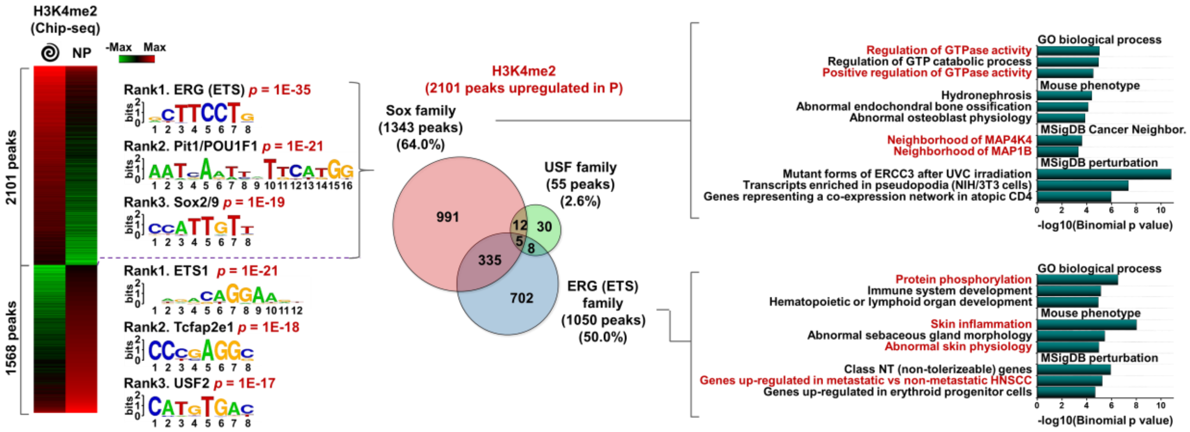


**Fig. S9.** Heatmap of H3K4me2 ChiP-seq signal for cells cultured on spiral geometry or non-patterned substrates. The top three predictive transcription factor motifs with p-values. Enriched annotations of genes for cells cultured on spiral patterns that contain a specific motif (SOX or *ETS* family) within the promoter. Venn diagram showing the number of enriched genes for cells cultured on spiral patterns linked to *SOX*, *ETS*, and *USF* families among H3K4me2-marked genes.

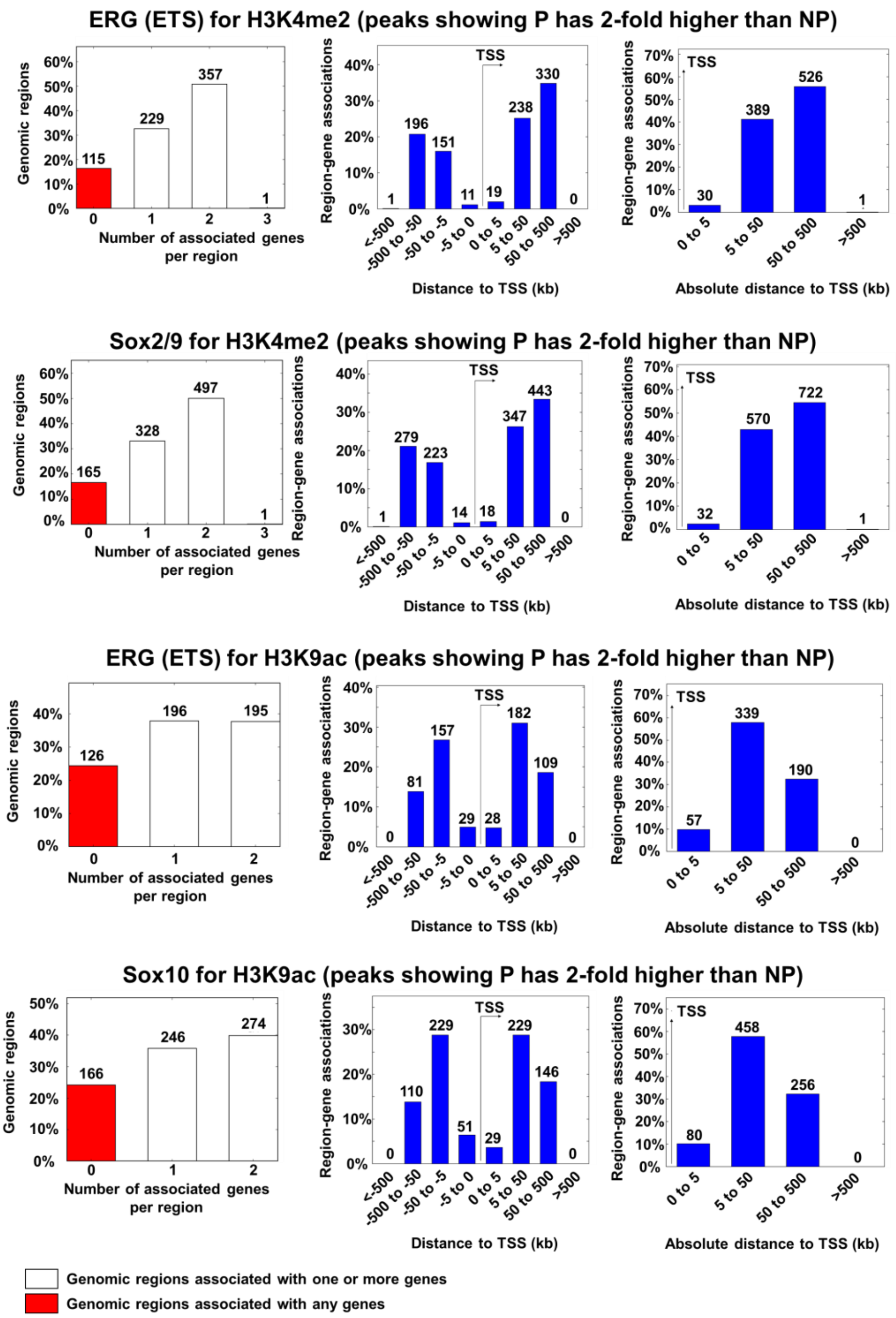


**Fig. S10.** Number of associated genes per region and binned by distance (with orientation or absolute value) to generate enriched annotations (GREAT) of genes for cells cultured on spiral patterns that contain a specific motif (*SOX* or *ETS* family) within the promoter.

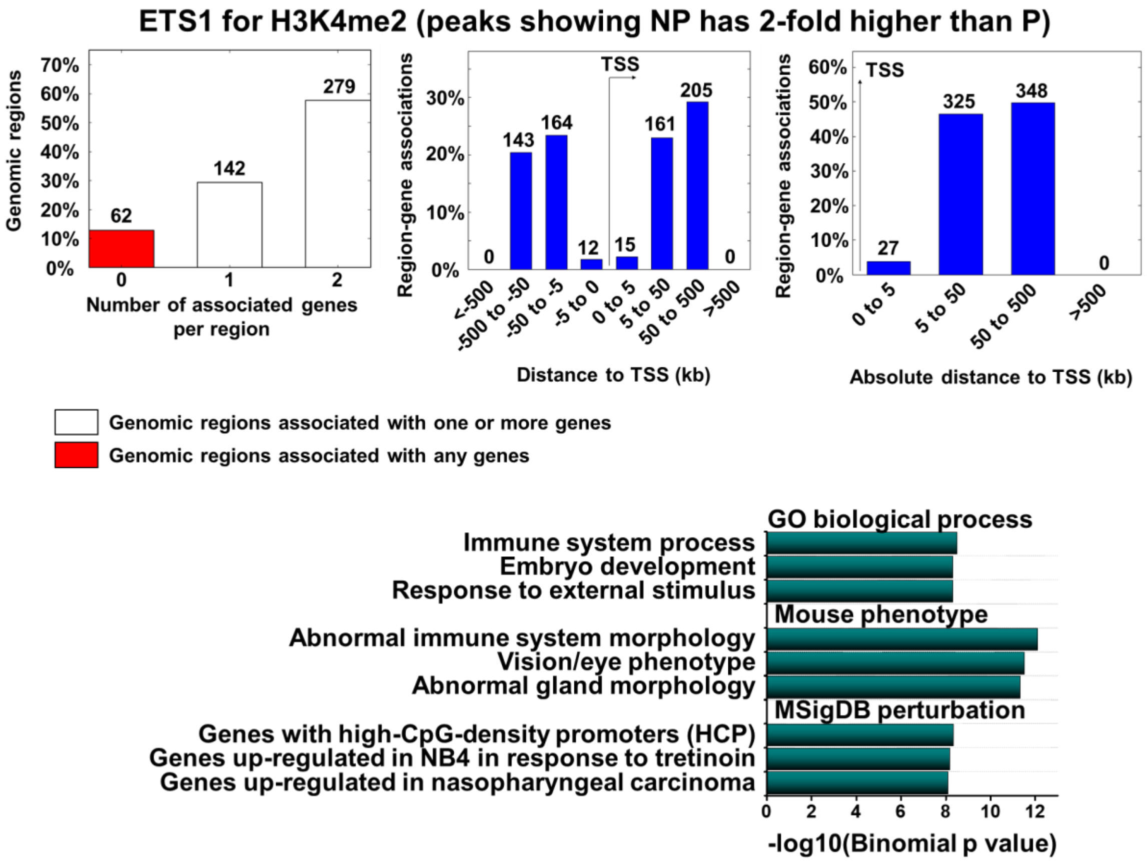


**Fig. S11.** Number of associated genes per region and binned by distance (with orientation or absolute value) to generate enriched annotations (GREAT) of genes for cells cultured on non-patterned substrates that contain a specific motif (*ETS1* family) within the promoter, and the enriched annotation results.

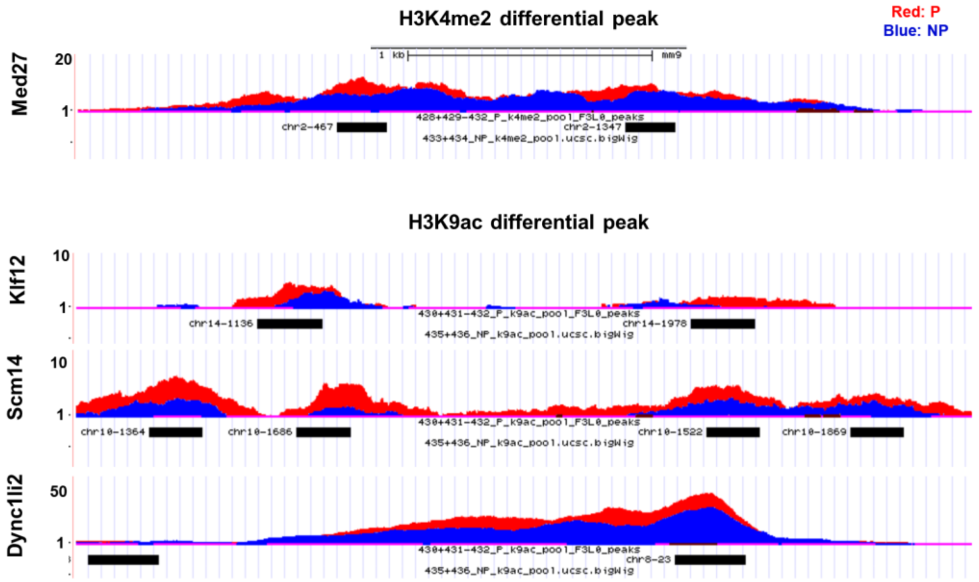


**Fig. S12.** Differential peaks of H3K4me2 (*Med27* and *Trim14*) and H3K9ac (*Klf12*, *Scml4*, and *Dync1li2*) associated with cells cultured on patterned gels compared to those cultured on non-patterned gels.

| Sample | # overlaps | p-value |
|---|---|---|
| P_k4me2 | 1758 | 2.74E-28 |
| P_k9ac | 870 | 9.72E-22 |
| NP_k4me2 | 1652 | 2.14E-25 |
| NP_k9ac | 917 | 5.18E-23 |
| PvsNP_k4me2 | 42 | 4.13E-01 |
| PvsNP_k9ac | 52 | 2.21E-02 |
| NPvsP_k4me2 | 40 | 2.28E-02 |
| NPvsP_k9ac | 61 | 1.06E-03 |

**Table S1.** Peak co-occurrence of *PRDM14* with the H3K9ac and H3K4me2 marks from our study within 1000bp.

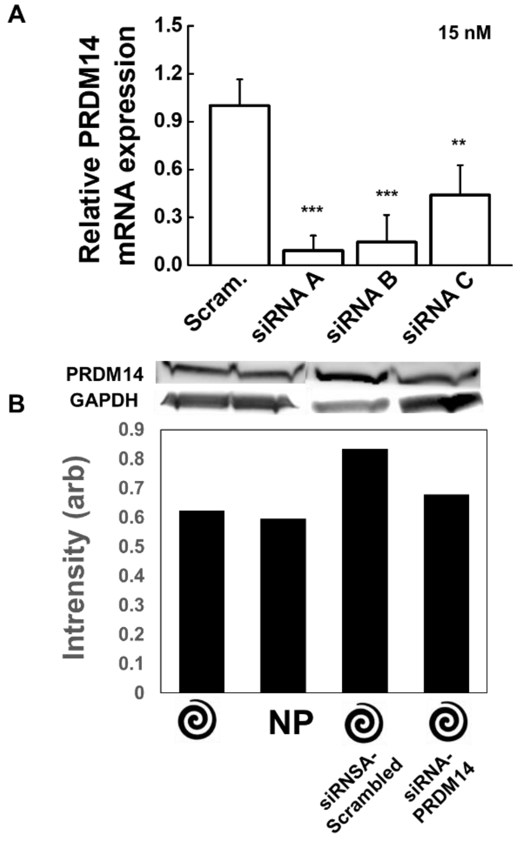


**Fig. S13.** (**A**) Results of real-time PCR to measure the gene expression of *PRDM14* (three different sequences of *PRDM14* siRNAs (A, B, and C) with 15 nM concentrations. (**B**) Western blots for *PRDM14* with scrambled or siRNA against *PRDM14*.

| Name | Company | Catalog # |
|---|---|---|
| High Glucose DMEM | Corning | 10-013-CV |
| Fetal bovine serum (FBS) | Denville | FB5001 |
| 12 well plates | VWR | 10062-894 |
| pen streptomycin | GIBCO | 15140122 |
| Trypsin 0.25% | GIBCO | 15050-065 |
| Sodium periodate | SIGMA-ALDRICH | 311448 |
| Fibronectin | SIGMA | F2006 |

**Table S2.** Reagent information for cell culture.

| Antibody | Company | Catalog # | Dilution/ Application |
|---|---|---|---|
| DAPI | INVITROGEN | D3571 | 1:5000/IF |
| Actin | INVITROGEN | A12379 | 1:200/IF |
| Goat 488-anti-rabbit | ABCAM | AB150077 | 1:200/IF |
| Goat 647-anti-moue | ABCAM | AB150115 | 1:200/IF |
| CD271 | ABGENT | AM1842a | 1:250/IF, FC |
| Jarid1B | BETHYL | A301-813A | 1:500/IF, FC |
| Oct4 | ABCAM | AB27985 | 1:500/IF, FC |
| Sox2 | ABCAM | AB97959 | 1:250/IF, FC |
| Nanog | SIGMA | N3038 | 1:500/IF, FC |
| Stat3 | ABCAM | AB119352 | 1:500/IF |
| a5b1 | MILLIPORE | MAB1969 | 1 µg/ml, blocking |
| H3K4me1 | CELL SIGNALING | 5326 | 1:250/IF, FC |
| H3K4me2 | CELL SIGNALING | 9725 | 1:250/IF, FC |
| H3K4me3 | CELL SIGNALING | 9751 | 1:250/IF, FC |
| H3K36me2 | CELL SIGNALING | 2901 | 1:250/IF, FC |
| H3K9me3 | CELL SIGNALING | 13969 | 1:250/IF, FC |
| HDAC1 | CELL SIGNALING | 5356 | 1:500/IF, FC |
| HDAC2 | CELL SIGNALING | 5113 | 1:500/IF, FC |
| HDAC3 | CELL SIGNALING | 3949 | 1:500/IF, FC |
| AcK | CELL SIGNALING | 9441 | 1:500/IF, FC |
| H3K4ac | ABCAM | AB113672 | 1:250/IF, FC |
| H3K9ac | CELL SIGNALING | 9649 | 1:250/IF, FC |

**Table S3.** Antibody information for immunostaining, flow cytometry analysis, and integrin blocking.

| Primary | Forward | Reverse |
|---|---|---|
| Jarid1B | GACATCACAAGCGAATGGTG | CGCTTTCATCCACAAGATCC |
| CD271 | GGG GGT AGA CCT TGT GAT CC | GTG TGC GAG GAC ACT GAG C |
| Oct4 | TGC CCG AAA CCC ACA CTG | CTC GGA CCA CAT CCT TCT CG |
| Sox2 | TGC TGC CTC TTT AAG ACT AGG AC | CGC CGC CGA TGA TTG TTA TT |
| Nanog | AAC AGG TGA AGA CCT GGT TCC | GAG GCC TTC TGC GTC ACA C |
| HDAC1 | CAA ATT GTG AGT CAT GCG GA | GGC ACC AAG AGG AAA GTC TG |
| GAPDH | TGC CTC GAT GGG TGG AGT | GCC CAA TAC GAC CAA ATC AGA |

**Table S4.** RT-PCR primer sequence information.
